# Extending the causal inference toolkit for ecologists: A visualization method of indirect effects

**DOI:** 10.64898/2026.02.09.704914

**Authors:** Allen Bush-Beaupré, Simon Coroller-Chouraki, Marc Bélisle

## Abstract

Much ecological research focuses on phenomena where a given variable can affect another either directly or indirectly through the effect of one or more intervening variables. While various methods to quantify the magnitude of these effects are available, they can be difficult to interpret in a meaningful way, especially for indirect effects. Studies thus often quantity direct effects and infer indirect ones more or less formally. We propose a method for visualizing indirect effects by means of plotting model predictions of an outcome in the presence of both exposure and mediator variables. We demonstrate the method through simulations and apply them to a real-world example involving Tree Swallows (*Tachycineta bicolor*) and their obligatory hematophagous ectoparasites, *Protocalliphora* bird blowflies (Diptera: Calliphoridae). Our procedure, which can be seamlessly integrated into an analyst’s workflow using commonplace software, should prove instrumental to disentangle and interpret relationships among variables involved in ecological mechanisms.

## Introduction

Ecological phenomena typically result from the interplay of several to numerous variables which can act on one another in three ways: directly, indirectly through the effect of one or more intervening variables, or both directly and indirectly. Importantly, the total effect of one variable on another is the sum of its direct and indirect effects, which can be positive, negative, or offset each other partially or completely. Identifying and quantifying direct and indirect effects is thus crucial to measure and interpret causal relationships to limit biases as much as possible (e.g., Cinelli et al., 2022). The resurgence of causal inference in several ecological fields has sparked interest in quantifying these effect types to understand the mechanisms underlying a wide range of phenomena (Arif & MacNeil, 2023; Dudney et al., 2024; Grace et al., 2025; Schrodt et al., 2025; Siegel & Dee, 2025). These include, among others, cascading effects in food webs (Buck, 2019; Bukovinszky et al., 2008), mutualistic relationships (Cosmo et al., 2023), and species interactions in the context of biological invasions (White et al., 2006). That said, whereas total and direct effects are (relatively) straightforward to assess, interpret, and report using regression model summaries and typical conditional prediction plots (e.g., Arel-Bundock, 2026; Fox & Weisberg, 2018), indirect effects may prove challenging to the point of hindering the progress of mechanistic research. Given the pervasive nature of indirect effects in ecology, it becomes imperative to develop methods that facilitate their quantification and visualization.

Modern causal investigations typically rely on Directed Acyclic Graphs (DAGs; Pearl et al., 2016; Shipley, 2016). Briefly, DAGs depict the hypothetical set of causal links among variables (observed or not) expected to play a role in a given phenomenon or system. As randomized controlled trials are generally not feasible in ecological research, the causal effect (total, direct, or indirect) of a variable on a specific response (outcome) variable is alternatively measured by statistically controlling for the influence of variables that may bias the relationship of primary interest. DAGs are instrumental for limiting biased effect measurements in that they dictate the specific set of variables that need to be statistically controlled for (Cinelli et al., 2022).

Several methods, such as path analysis or structural equation modelling, have been developed to dissect and quantify causal links within DAGs (Shipley, 2016). Omitting details, these methods generally consist of fitting regression models for each (outcome) variable which is hypothesized to be caused by other (exposure) variables (see Pearl et al. (2016) and references therein for detailed workflows and limitations. Basically, the direct causal effect of a given exposure variable on an outcome variable is estimated by its regression coefficient when controlling for (1) at least one intervening (mediator) variables involved in each of its indirect effects on the outcome variable, and (2) variables that may generate spurious associations through backdoor paths (i.e., by applying a back or front-door adjustment). Estimating the total causal effect of an exposure variable commands adjusting for the same set of variables as above but excluding mediators and their causal descendants (other than the outcome) if present. Finally, estimating the indirect effect of an exposure variable on an outcome variable through a mediator consists of three steps. First, the mediator is regressed on the exposure. Second, the outcome is regressed on the mediator. If it is hypothesized that the exposure also has a direct effect on the outcome, that variable is also included in the second regression. Third, the indirect effect of the exposure on the outcome through the mediator variable is estimated by multiplying the regression coefficient of exposure on mediator with the regression coefficient of mediator on the outcome. However, this simple procedure is only applicable in cases where, among other things, there are no backdoor paths, the relationships are linear, and there is no interaction between the exposure and the mediator (Pearl, 2014; Vanderweele, 2015). The last two constraints are quite limiting outside the domain of applicability of Gaussian-distributed variables and linear models. Indeed, intrinsically nonlinear relationships, interactions, and ecological measures such as counts and proportions that require different distributional assumptions in the context of Generalized Linear Models (GLMs), are legion in ecology. Moreover, even when GLMs are linear and do not involve interactions on the scale of the link function, they can remain nonlinear and include inherent interactions on the response scale (e.g., with log or logit link; Spake et al., 2023). Lastly, although some counterfactual-based mediation techniques allow to compute indirect causal effects under such conditions (Pearl, 2014; Vanderweele, 2015), their application, interpretation, and visualization remain unintuitive to most practitioners.

We propose a simple intuitive procedure for visualizing the indirect effects of a variable of interest through a mediator. After demonstrating its use through simulation, we apply the procedure to a real-world ecological study system and dataset involving Tree Swallows (*Tachycineta bicolor*) and obligatory hematophagous ectoparasites of their nestlings, namely *Protocalliphora* spp. bird blowflies (Diptera: Calliphoridae). Specifically, we assess how Tree Swallow egg hatching date affects the abundance of bird blowflies in nests both directly and indirectly through a proxy of host availability (nestling-days) and the temperature to which bird blowfly larvae were exposed.

### The Procedure with Simulated Data

Generalized linear models, model predictions, and visualizations for the simulation described below were conducted using *glmmTMB* (v.1.1.13; Brooks et al., 2017), *marginaleffects* (v.0.30.0; Arel-Bundock et al., 2024), and *ggplot2* (v.4.0.0; Wickham, 2016) packages, respectively, in the R environment (v.4.5.1; R Core Team, 2025).

We illustrate the visualization procedure for the simple mediation phenomenon depicted in the DAG of Figure 1A. It relies on fitting two regression models – one for the direct effect of the exposure variable (E) on the mediator (M) (M ∼ E) and another for the direct effects of variables E and M on the outcome variable (O) (O ∼ E + M). We assumed the following. Variable E is normally distributed with mean 0 and standard deviation 1. Variable M is Poisson distributed with a rate *λ*_*M*_ caused solely by E with a slope (*β*_*EM*_) of 0.1 and an intercept (*α*_*M*_) of 4 on the log scale. Variable O is Bernoulli distributed with probability *ρ*_*O*_ caused by both E and M with slopes *β*_*OE*_ = 0.1 and *β*_*ME*_ = 0.05, respectively, and intercept *α*_*O*_ = -3 on the logit scale (Equation 1).

**Figure 1.**
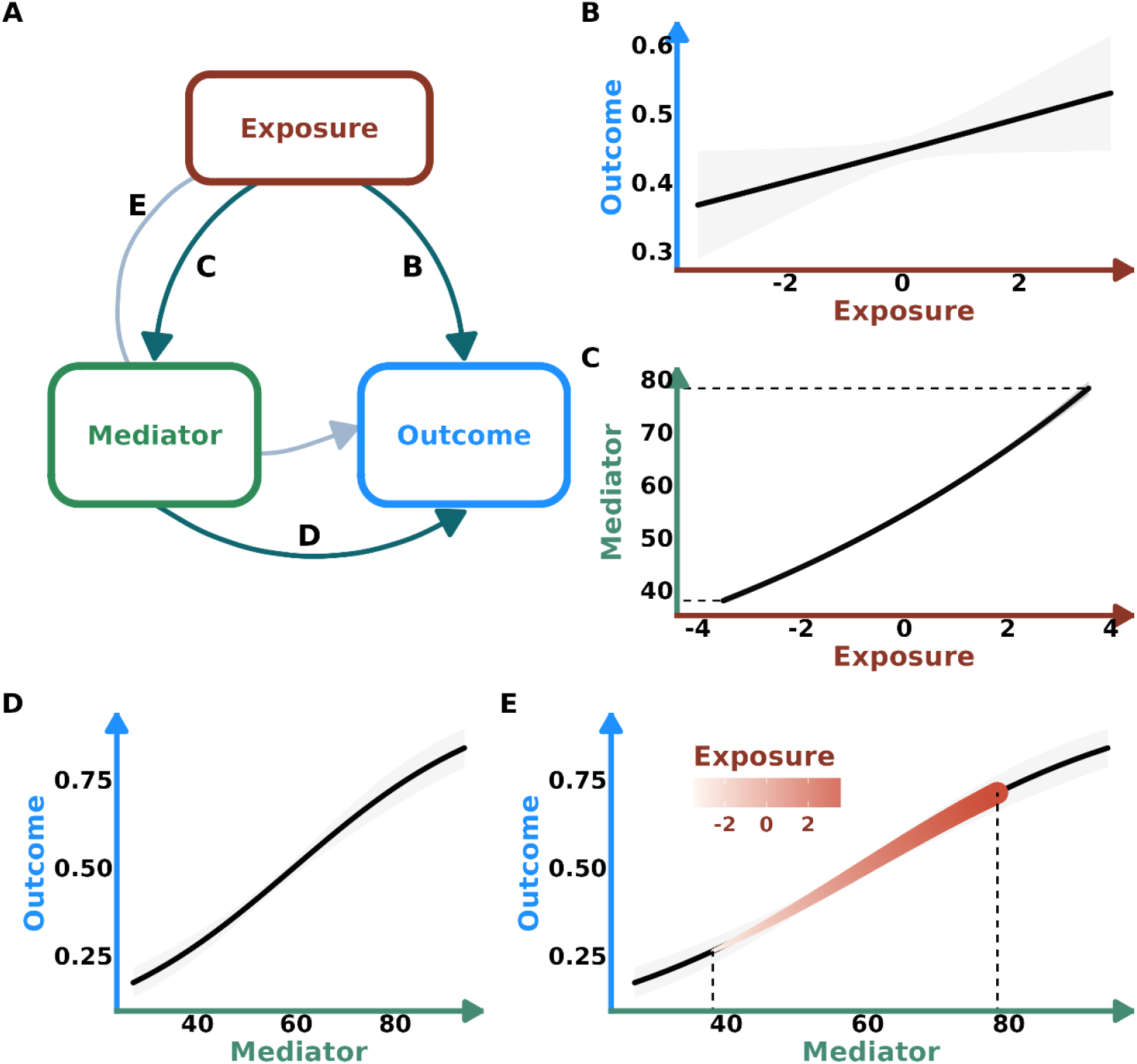
(A) Directed acyclic graph of a simulated example of the direct effect of an exposure variable on an outcome variable and its indirect effect through a mediator. (B) Prediction of the direct effect of the exposure on the outcome. (C) Prediction of the direct effect of the exposure on the mediator. (D) Prediction of the direct effect of the mediator on the outcome. (E) Prediction of the indirect effect of the exposure on the outcome superimposed on the prediction of the direct effect of the mediator on the outcome. Predictions generated from Generalized Linear Models.

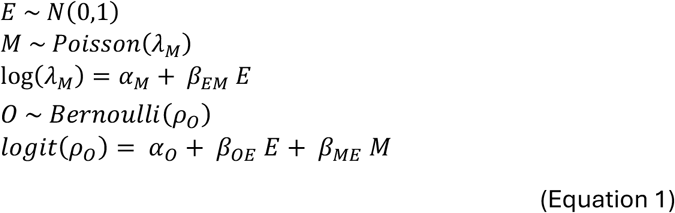

Visualizing direct effects is straightforward. One simply plots the expectation of the response variable of interest as a function of the predictor of interest while holding constant the value of the other covariate, if any (Figure 1B-D). These are the typical plots reported by authors. On the other hand, visualizing the indirect effect of E on O through M (Figure 1E) relies on fitting a GLM according to Equation 1 (O ∼ E + M) and generating expected values of O for the expected values of M as a function of E (E[O] ∼ E + E[M], where E[M] ∼ E). One can then display the indirect effect of E on O through M as a superposition of the expected values of O as a function of the full range of values of M (E[O] ∼ M; Figure 1D) and the expected values of O as a function of the expected values of M predicted by E (E[O] ∼ E[M]; Figure 1E). As the expected values of M as a function of E (section between dotted lines in Figure 1C) do not cover the entire range of possible M values, the expected values of O as a function of the expected values of M only represent a portion of the full relationship between M and expected values of O (section between dotted lines in Figure 1E). This section of the relationship represents the indirect effect of E on O through M and is displayed here by changing the color and width of the trend line proportional to E. See code for additional details.

In the case of this simulation, as E has a moderate effect on O (∼15-percentage-point (%pt.) increase; Figure 1B), a strong effect on M (∼100% increase; Figure 1C) and M has a strong effect on O (∼60%pt. increase; Figure 1D), the indirect effect of E on O through its effect on M (∼50%pt. increase; Figure 1E) turns out to be stronger than its direct effect. The absence of such an accessible method for visualizing indirect effects likely contributes to the tendency of researchers to focus solely on direct effects. This, coupled with the tendency of “overcontrolling” for potential confounders (Cinelli et al. 2022), can thus in turn lead to interpretations in which the influence of certain variables (e.g., variable E in the above simulation) is erroneously deemed unimportant.

### Case Study: *Protocalliphora* Abundance in Swallow Nests

The larvae of *Protocalliphora* spp. bird blowflies are among the most prevalent ectoparasites of altricial bird nestlings across the Holarctic region (Bennett & Whitworth, 1991; Sabrosky et al., 1989). Despite a vast and debated literature on the impact of these obligatory hematophagous ectoparasites on their hosts’ fitness, few studies have addressed the determinants of the abundance of *Protocalliphora* larvae within host nests (Coroller-Chouraki et al., 2026a,b). While Coroller-Chouraki et al. (2026b) examined the effects of several putative causes of the abundance of *Protocalliphora* spp. in the nests of Tree Swallows breeding along a gradient of agricultural intensity, they focused on direct effects and neglected to formally quantify or visualize hypothesized indirect effects. Here, we apply our novel visualization procedure to two indirect effects linked to the reproductive timing of swallows and included in the DAG on which the aforementioned study was based. Specifically, we address the mediating effect of nestling availability (Figure 2) and that of the temperature experienced during the larval stage (Figure 3) on *Protocalliphora* spp. abundance. In the first case (Figure 2), it was hypothesized that the hatching time of swallow eggs will not only influence *Protocalliphora* spp. abundance through the temporal matching between the respective phenology of the blowfly and its bird host (direct effect), but also because swallows that breed early in the season are expected to be in better body condition and thus to produce larger clutches and broods with higher survival rates (Forslund & Pärt, 1995; Nooker et al., 2005), which would provide greater nestling (resource) availability for blowfly larvae to feed upon (Hurtrez-Boussès et al., 1999). To isolate these specific hypothesized effects, we had to control for backdoor paths linked to the variation in agricultural intensity, weather conditions, and female swallow age associated to hatching date, nestling availability, and/or *Protocalliphora* spp. abundance through different mechanisms (e.g., insect prey availability to swallows, female experience; see Coroller-Chouraki et al. 2026b; Table 1). The second case (Figure 3) looks at the mediating effect of temperature during *Protocalliphora* larval development. Tree Swallows breed in late spring and early summer when temperatures are rising. The rate of growth and development of *Protocalliphora* spp. larvae have been found to increase with temperature, suggesting that this variable partly determines the quality of the growing and developmental conditions experienced by the larvae, and thus their survival and number in the nest (Bennett & Whitworth, 1991; Dawson et al., 2005; Mennerat et al., 2021). As for the mediating effect of nestling availability, backdoor paths had to be blocked by controlling for specific variables (Table 1).

**Table 1.** Models used to quantify the mediating effects of host availability and temperature during the larval development on *Protocalliphora* spp. abundance within infested Tree Swallow nests monitored on 40 farms from Southern Québec between 2004 and 2019. Models follow the causal inference framework found in Coroller-Chouraki et al. (2026b).

| Outcome variable | Distribution* (model type and link function) | Exposure, mediator, and control variables ** |
| --- | --- | --- |
| Host availability (nestling-days) | Gaussian (GAMM with log link) | s[hatching date] + [agricultural landscape] + [mother age] + [temp<sub>S</sub>*precip<sub>S</sub>] + [temp<sub>N</sub>*precip<sub>N</sub>] |
| Temperature during *Protocalliphora* development | Lognormal (GLMM with identity link) | [hatching date] |
| *Protocalliphora* abundance | Tweedie (GAMM with log link) | s[hatching date] + [nestling-days] + te[temp<sub>P</sub>*precip<sub>P</sub>] + [agricultural landscape] + [temp<sub>S</sub>*precip<sub>S</sub>] |

\* The Generalized Additive Mixed Models (GAMM) and Generalized Linear Mixed Model (GLMM) were fitted with the *mgcv* (v.1.8.42; Wood, 2017) and *glmmTMB* (v.1.1.13, Brooks et al., 2017), respectively. All models included random intercepts for farm ID nested within year. s() and te() refer to smoothing terms and tensor products, respectively.
\*\* temp<sub>□</sub> & precip<sub>□</sub>: temperature and precipitation during early spring (subscript S), during nestling development (subscript N), and during *Protocalliphora* development (subscript P)

**Figure 2.**
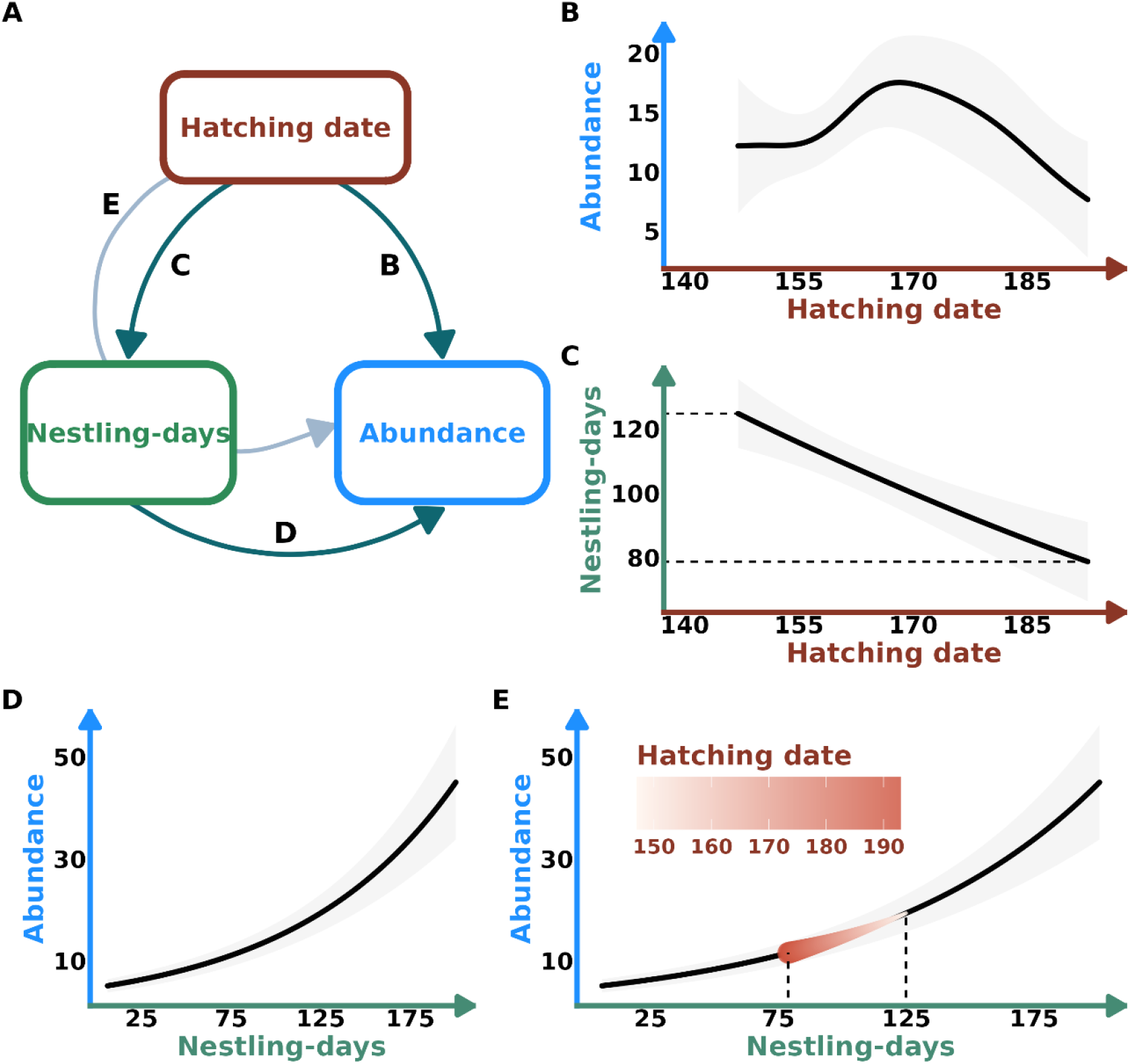
(A) Directed acyclic graph of the direct effect of first hatching date on *Protocalliphora* abundance and its indirect effect through nestling-days. (B) Prediction of the direct effect of first hatching date on *Protocalliphora* abundance. (C) Prediction of the direct effect of the first hatching date on nestling-days. (D) Prediction of the direct effect of nestling-days on *Protocalliphora* abundance. (E) Prediction of the indirect effect of first hatching date on *Protocalliphora* abundance superimposed on the predictions of the direct effect of nestling-days on *Protocalliphora* abundance. Predictions generated from Generalized Additive Mixed Models.

**Figure 3.**
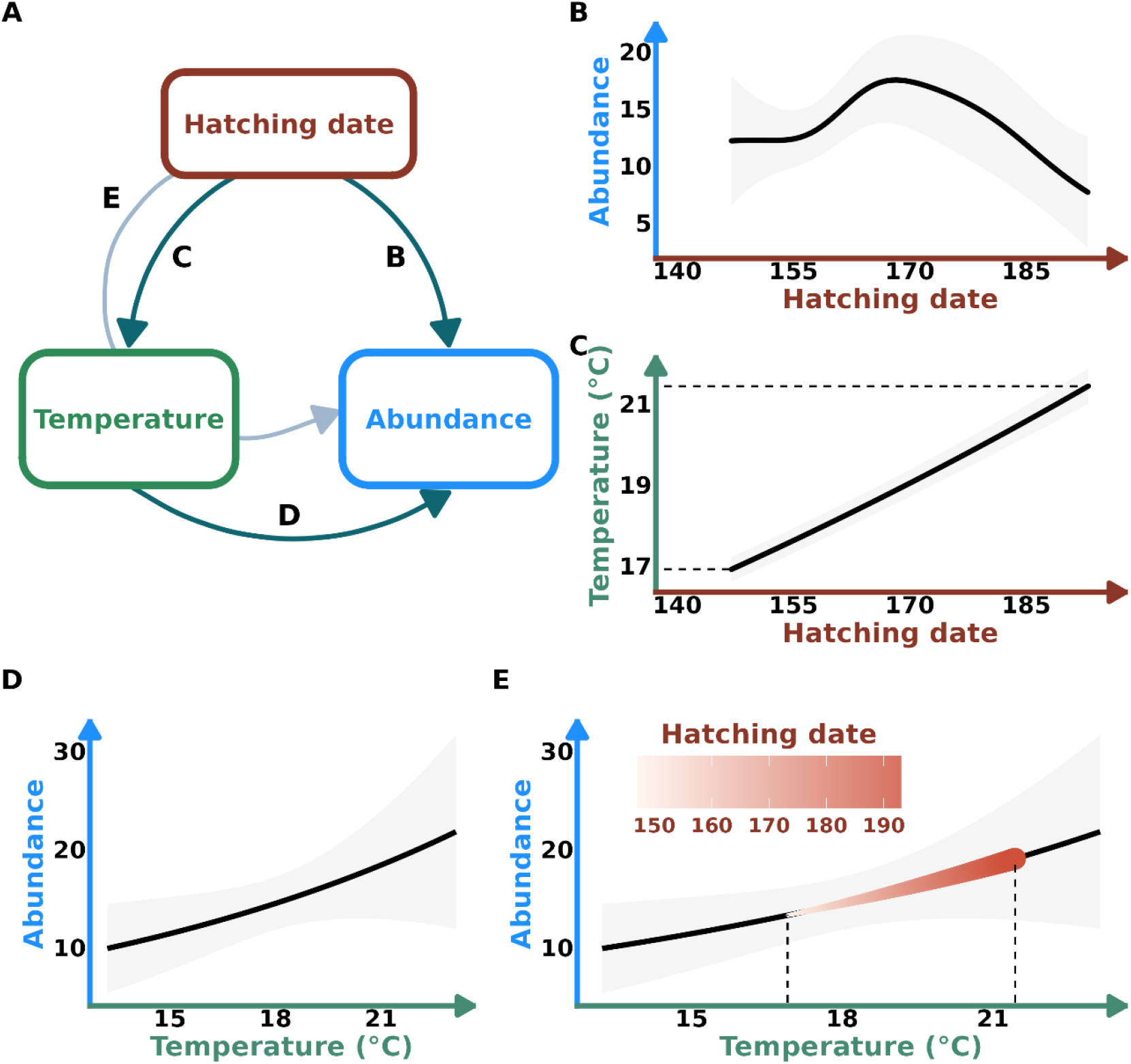
(A) Directed acyclic graph of the direct effect of first hatching date on *Protocalliphora* abundance and its indirect effect through temperature. (B) Prediction of the direct effect of first hatching date on *Protocalliphora* abundance. (C) Prediction of the direct effect of the first hatching date on temperature. (D) Prediction of the direct effect of temperature on *Protocalliphora* abundance. (E) Prediction of the indirect effect of first hatching date on *Protocalliphora* abundance superimposed on the predictions of the direct effect of temperature on *Protocalliphora* abundance. Predictions generated from Generalized Additive Mixed Models.

The data used to fit the models required to estimate the direct and indirect effects of interest come from a study on the breeding ecology of Tree Swallows within a network of 40 farms, each with 10 nestboxes, distributed across a 10,200-km^2^ gradient of agricultural intensity in Southern Québec, Canada (see Garrett et al., 2022; Ghilain & Bélisle, 2008). Nestboxes were monitored every two days during the breeding seasons of 2004 through 2019 to gather data on occupancy; species, age, sex, and ID of occupants; clutch and brood sizes; laying, hatching, and fledging dates; as well as nestling growth, survival, and fledging success. At the end of the season, all nests were collected to count the *Protocalliphora* spp. puparia therein to estimate *Protocalliphora* abundance during the nestling phase. A total of 1,175 nests infested by *Protocalliphora* spp. were processed across the 16-year study. We estimated host availability for *Protocalliphora* larvae by summing the number of days during which each nestling remained alive until it fledged (thereafter referred to as nestling-days). *Protocalliphora* larvae typically initiate their development at the onset of the nestling phase and pupate seven days after nestlings have fledged (Sabrosky et al., 1989). We hence averaged the hourly temperature (and precipitation) data for each swallow brood during its nestling phase extended by seven days. Note that Coroller-Chouraki et al. (2026b) considered that the effect of temperature during this extended time window on *Protocalliphora* abundance may depend on the amount of precipitation but failed to detect any such meaningful interaction. We thus chose to model the effect of hatching date (of first egg) only on temperature (Temperature model) and assess its indirect effect on *Protocalliphora* abundance while setting precipitation at its mean value (Table 1). The Generalized Additive Mixed Models (GAMM) and Generalized Linear Mixed Model (GLMM) were fitted with the *mgcv* (v.1.8.42; Wood, 2017) and glmmTMB (v.1.1.13, Brooks et al., 2017), respectively.

The visual representations of the direct effect of hatching date on *Protocalliphora* abundance and its indirect effects through nestling-days and temperature are displayed in Figures 2 and 3, respectively. *Protocalliphora* abundance showed a near quadratic relationship with the timing of the first hatching event of swallow broods, peaking at 17.7 individuals (95% CI = [13.6, 21.7]) for broods that started to hatch on day of year 168 (Fig. 2B & 3B). However, host availability in terms of nestling-days decreased (linearly) by approximately 33% with hatching date (Fig. 2C). *Protocalliphora* abundance increased (exponentially) nearly 250% across the range of nestling-day values (Fig. 2D). The resulting indirect effect of hatching date on *Protocalliphora* abundance through nestling-days was thus negative due to the lower host availability provided by later broods; an indirect effect size of nearly -50%. This effect is displayed by the gradient of hatching date that increases in the opposite direction than the positive trend caused by nestling-days (Fig. 2E). Regarding the temperature to which *Protocalliphora* were exposed, it increased by ∼30% across the range of hatching dates (Fig. 3C). As temperature increased across its observed range, *Protocalliphora* abundance increased by ∼75% (Fig. 3D). As a result, hatching date increased *Protocalliphora* abundance indirectly through temperature by ∼50% (Fig. 3E).

## Discussion

Indirect effects are omnipresent in ecological phenomena. Although much effort has been focused on developing methodological frameworks by which the magnitude of indirect effects may be quantified (e.g., VenderWeele 2015), the use and interpretation of such measures remain complex and uncommon in the ecological literature. Using a simulated example and case study, we demonstrate a straightforward method to visualize indirect effects of a variable of interest across a mediator. We believe it can serve as supplemental means to convey the information about indirect effects in a more meaningful way compared to reporting coefficient values from regression outputs or within a DAG as is typically done in path analysis or structural equation modeling (e.g., Shipley 2016). While simple, our method should prove instrumental to visualize and interpret relationships that are inherently difficult to comprehend.

The visual representation of direct and indirect effects displayed here provides a more intuitive means to decompose and interpret effect types. For instance, in the first example from our case study, the visualization of the indirect effect of Tree Swallow hatching date on *Protocalliphora* abundance through its own negative effect on host availability (Figure 2C) led to a 50% decrease across the range of observed hatching dates, lending support to the hypothesis that early nesters are better breeders (Forslund and Pärt 1995; Figure 2E). This contrasts with the 35% decrease in *Protocalliphora* abundance across the range of observed hatching dates (Figure 2B). In the second example, the visualization clearly showed that a late hatching date increases the temperature to which the ectoparasites are exposed (Figure 3C), and that this, in turn, increases *Protocalliphora* abundance (Figure 3E). This is consistent with the result that bird blowfly larvae tend to survive better in experimentally-heated nests of Tree Swallows (Dawson et al., 2005).

Considering direct and indirect effects as above also reduces the likelihood of misinterpretation, since opposing effects may cancel each other out, or mediators may mask the effect of a causal ancestor (Shipley, 2016). Selecting variables through stepwise procedures or by comparing models based on an information criterion but without accounting for the causal structure underlying the phenomenon of interest, is prone to produce such misinterpretation (Cinelli et al., 2022). Studies by Mennerat et al. (2021) and Maziarz et al. (2022) exemplify this as hatching date has been discarded through such variable selection procedure, casting doubts as to its role in determining *Protocalliphora* spp. abundance through its effect on temperature in Blue Tit (*Cyanistes caeruleus*) and Wood Warbler (*Phylloscopus sibilatrix*) nests, respectively. Besides, incorporating a mediator in a composite exposure or outcome variable like a ratio can also complicate interpretation and bias effect sizes (Kronmal, 1993; Mooldijk et al., 2025). Such a situation can be found in the work of Mennerat et al. (2021) who modeled the number of *Protocalliphora* spp. larvae per nestling as a function of hatching date and temperature; variables that can be related directly and indirectly to both the numerator and denominator of the ratio. In a causal inference context (in contrast to a predictive one), including the ratio denominator (i.e., bird brood size) as an offset would not have fixed the problem as its slope parameter would have been fixed to one and thus not free to vary. Enabling straightforward visualization of indirect effects would likely encourage clearer specification of causal links among study variables and thereby help researchers avoid the above pitfalls. That said, we emphasize that all causal inference methods ultimately depend on the validity of the underlying DAG to which they are applied, including the assumption that no unobserved (or unknown) variables create backdoor paths.

On another front, although counterfactual-based mediation techniques allow to compute indirect causal effects under a wide diversity of data types and functional structure (Pearl, 2014; Vanderweele, 2015), they are nevertheless often only accessible to insiders. Hence, focusing on visual representations of model predictions and effect sizes as done here opens the causal inference framework to a wider community of practice. Moreover, our visualization method is not restricted to the use cases presented here as it can be adapted to cases where the exposure variable does not have a direct effect on the outcome of interest, such as the indirect effect of a predator on producers through its prey. The method may also prove very useful to complement studies on the direct (e.g., toxicological) and indirect (e.g., trophic) effects of contaminants such as pesticides on ecological systems (e.g., Fleeger, 2020; Saaristo et al., 2018). Effect visualizations may indeed facilitate a deeper understanding of such complex relationships among members of the public and stakeholders, thereby increasing the likelihood of informed and appropriate actions. More complex indirect effects involving interactions and multiple mediators of the carry-over effects of snow characteristics on Bighorn Sheep (*Ovis canadensis*) life parameters have been visualized in an additional case study found at our accompanying website. Further exemplifying the scope of our method, Briau et al. (2025) used it to illustrate the indirect effect of the daily number of visits to feeders on estrus probability mediated by spring body mass in a wild Eastern Chipmunk (*Tamias striatus*) population. While the authors did not interpret the indirect effect in and of itself, our method allowed for a clear visual representation of its considerable magnitude. As the visualization method becomes more frequently used, it is our hope that the interpretations made from the results of such complex analyses become richer, adding nuances and being more intuitive for readers.

## Acknowledgements

The *Protocalliphora* data come from a long-term study of Tree Swallow breeding ecology conducted by and funded through grants to MB, Fanie Pelletier, and Dany Garant from U de Sherbrooke. We are indebted to all farm owners and the numerous graduate students and research assistants who helped collect and process samples and data. This work was conducted under the approval of the animal care committee of U de Sherbrooke and was financially supported by Natural Sciences and Engineering Research Council of Canada discovery grants to FP, DG, and MB, team research grants from the Fonds de recherche du Québec—Nature et technologies to FP, DG,and MB, by the Canada Research Chairs program to FP and MB, as well as by the Canadian Foundation for Innovation to FP, DG,and MB, G&Cs from the Canadian Wildlife Service of Environment and Climate Change Canada and the Université de Sherbrooke. We also thank François Briau for contributing to the aesthetic of the visualized indirect effects.

## Data availability

All code and data used for the simulations and data analysis can be found at 10.5281/zenodo.18564136. A tutorial demonstrating the simulated cases and case study can be found at https://abushbeaupre.github.io/indirect_viz/

